# NLRP3 inflammasome-related microglial pyroptosis in EcoHIV infected mice

**DOI:** 10.64898/2026.04.29.721781

**Authors:** Hailong Li, Charles F Mactutus, Diego Altomare, Michael Shtutman, Rosemarie M. Booze

**Affiliations:** Department of Psychology, Cognitive and Neural Science Program, University of South Carolina, Columbia, South Carolina, USA; Department of Drug Discovery and Biomedical Sciences, College of Pharmacy, University of South Carolina, Columbia, USA

**Keywords:** NLRP3, Microglia, Inflammation, Pyroptosis, EcoHIV

## Abstract

HIV-associated neurocognitive disorders (HAND) have become a major clinical concern, particularly among the aging HIV-1-seropositive population, which is generally characterized by persistent viral reservoirs and a lower level of chronic inflammation. NLRP3 inflammasome activation exhibits its unique role in the progression of many chronic inflammatory diseases. Furthermore, pyroptosis, an inflammatory form of programmed cell death, has been implicated in numerous neurological diseases. However, the mechanisms linking EcoHIV infection, microglial pyroptosis, and NLRP3 inflammasome activation remain incompletely understood. In this study, EcoHIV was retro-orbitally injected into C57BL/6J wild-type mice and analyzed at 14-, 30-, 60-, and 90-days post-infection to establish a NeuroHIV model. Additionally, *in vitro*, BV2 microglial cell line was infected with EcoHIV and treated with MCC950, an inhibitor of the NLRP3 inflammasome, for three days. Pyroptosis marker GSDMD, NLRP3 inflammasome components, Caspase-1 (a marker of inflammasome activation), HLA-DR (an immune activation marker), Programmed-death 1 (PD-1, an immune checkpoint molecule), and Ki67 (a cellular proliferation marker) were assessed by immunofluorescence staining. Results showed that EcoHIV-infected mice showed a peak in NLRP3 expression at 14 days post-infection, compared with controls, followed by a modest decline at 30 days, while GSDMD expression increased progressively across 14 and 30 days. These findings demonstrate dynamic changes in microglial pyroptosis and NLRP3 inflammasome activation over the course of EcoHIV infection. *In vitro*, EcoHIV-infected BV2 cells exhibited significantly increased EcoHIV-eGFP fluorescence compared with controls, confirming the utility of BV2 cells as an *in vitro* model of microglial EcoHIV infection. Expression levels of GSDMD and NLRP3 were elevated following infection, indicating enhanced pyroptosis and neuroinflammation. Treatment with MCC950 significantly reduced the expression of GSDMD, NLRP3, HLA-DR, PD-1, and Ki67, suggesting that inhibition of NLRP3 inflammasome activity suppresses both pyroptosis and microglial activation and proliferation. Together, elucidating the interplay between microglial pyroptosis and NLRP3 inflammasome activation may provide new insights into the pathogenesis and potential therapeutic strategies for NeuroHIV in the aging HIV-1-seropositive population.

## 1. Introduction

Human immunodeficiency virus type 1 (HIV-1) infection is characterized by a progressive loss of CD4⁺ T cells. Although chronic antiretroviral therapy (cART) effectively suppresses viral replication, HIV-1-associated neurocognitive disorders (HAND) continue to affect up to 60% of people living with HIV-1 [1, 2]. Persistent chronic inflammation within the central nervous system (CNS) is widely recognized as a key pathological feature of HAND. Previous studies using gp120 transgenic mice, which exhibit neuropathological and behavioral abnormalities characteristic of HAND, have demonstrated that chronic neuroinflammation plays a predominant role in gp120-induced neuropathology [3, 4].

Subsequent investigations revealed that approximately 95% of quiescent lymphoid CD4⁺ T cells undergo caspase-1-mediated pyroptosis during HIV-1 infection [5, 6]. Using an *ex vivo* human lymphoid aggregate culture (HLAC) system, HIV infection was shown to trigger this highly inflammatory form of programmed cell death, accompanied by the release of cytoplasmic contents, including pro-inflammatory cytokines, into the extracellular space [7, 8, 9]. Caspases, a family of cysteine proteases, are central mediators of regulated cell death pathways, including caspase-1-dependent pyroptosis [10, 11, 12]. Notably, among inflammasome sensors, NLRP3 has been identified as the primary driver of caspase-1 activation in CD4⁺ T cells during chronic HIV-1 infection. NLRP3-associated pyroptosis has been shown to constitute a dominant mechanism contributing to CD4⁺ T-cell depletion in patients with HIV-1 [13].

In addition to lymphocytes, exposure of monocytes to the HIV-1 neurotoxic protein Tat has been shown to stimulate interleukin-1β (IL-1β) secretion [14, 15]. Importantly, the persistence of Tat and other viral proteins in the brain is thought to sustain low-level neuroinflammation associated with HAND, even under cART [16, 17]. Microglia, which serve as major target cells for HIV-1 infection and act as viral reservoirs within the CNS, are capable of releasing Tat as well as numerous inflammatory mediators. However, the relationship between microglial pyroptosis and NLRP3 inflammasome activation during HIV-1 infection remains poorly defined.

Characterizing the spatial distribution of NLRP3 inflammasome-associated microglial pyroptosis across the entire brain is critical for understanding how inflammatory processes disrupt neural circuits and contribute to behavioral and pathological outcomes. In this study, we employed the BrightSLICE light-sheet microscopy platform in combination with the NeuroInfo system, providing an end-to-end solution for whole-brain imaging and quantitative analysis of NLRP3 inflammasome activation and microglial pyroptosis during EcoHIV infection. Our results demonstrate a critical role for NLRP3 inflammasome-dependent microglial pyroptosis in driving microglial dysfunction throughout HIV-1 infection, as evidenced by both *in vivo* and *in vitro* analyses.

## 2. Materials and Methods

### 2.1. EcoHIV virus infection

*C57BL6/J* mice were housed in AAALAC-accredited facilities according to guidelines established by the National Institutes of Health. All procedures were approved by the Institutional Animal Care and Use Committee (IACUC) of the University of South Carolina (Federal Assurance #D16-00028). Twelve mice were randomly assigned to receive retro-orbital injections of EcoHIV (n=13) or saline (control; n=6) in a controlled environment. Animals were maintained under a 12:12 light/dark cycle with *ad libitum* access to food and water. The detailed protocol about EcoHIV virus construction and retro-orbital injection were explained in our previous study [18]. BV2 cell line was purchased from ATCC (Cat. No. CRL-2469, ATCC) and cultured in DMEM/F12 growth medium with 10% fetal serum at 37°C, 5% CO^2^ condition. BV2 cells were cultured in the 12-well plate with glass bottom till 30% confluence. There are four groups: Control group; EcoHIV group (6μL, 1.26 × 10^6^ TU/mL); EcoHIV + MCC950 group (6μL of EcoHIV virus and 10μM of MCC950); MCC950 only group (10μM).

### 2.2. SLICE – Light sheet imaging

Mice were perfused with 4% paraformaldehyde (add 10% sucrose) and brain tissue was incubated with 4% paraformaldehyde at 4°C for overnight. Whole brain tissue was incubated in 30% sucrose in 1× PBS at 4°C until the sample sinks. For tissue clearing, brain tissue was incubated in the Binaree Tissue Clearing Rapid Solution (Cat. No. BRTC-402, BINAREE) at 37°C for 8 hours. Then sample was sequentially blotted with permeabilization solution (3 days), primary antibody (3 days, see Supplementary Table 1), secondary antibody (3 days) in a shaking incubator at 37°C. Brain samples will be incubated with mounting medium (Cat. No. BRMO-060, BINAREE) in a shaking incubator at 24°C for 3 days before imaging. For SLICE – Light Sheet Imaging, briefly, whole mouse brain was fixed in a 10×10 cuvette filled with the above mounting medium (Reflection index: 1.4587) and the fluorescent signals of samples were excited through the bi-lateral lasers with 10× objectors. Images were acquired via BrightSlice software and anatomic assessment was performed utilizing NeuroInfo software (MicroBrightfield, Williston, VT, USA).

### 2.3 Image volume acquisition & post-processing of BrightSLICE light sheet microscope

Brain-wide cell or signal detection were performed and displayed within the NeuroInfo software (Figure 1). A BINAREE-cleared mouse brain was registered to the Allen Mouse Brain Common Coordinate Framework (CCF) to generate a model of the equivalent position of the experimental slice on the reference atlas (Figure 1C,D). Region-specific quantitative fluorescence intensity was analyzed using NeuroInfo software, mPFC area (Figure 1C) mapped from the Allen CCF to the experimental volume. Next, fluorescence count and strength per Allen Mouse brain region separated by hemisphere were extracted across the entire registered brain.

**Figure 1.**
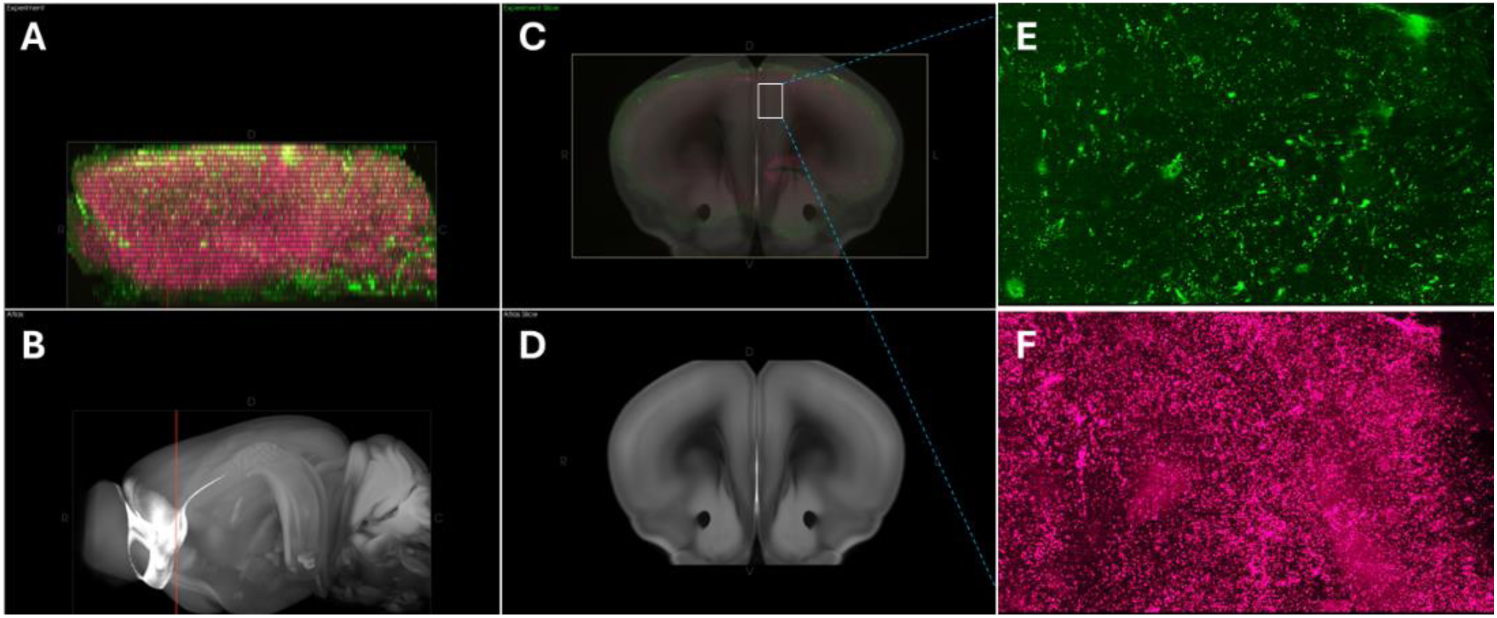
Whole brain image volume acquisition & post-processing via BrightSLICE light sheet microscope. Mouse brain was fixed with 4% PFA, cleared by BINAREE kit and imaged with SLICE light sheet microscope using a 10× objective. The BINAREE-cleared mouse brain was registered to the Allen Mouse Brain Common Coordinate Framework (CCF). (A) Maximum intensity projection of the whole brain with coronal red intersection line highlighting the location of the experimental image slice displayed in C. (B) The red plane indicates the equivalent position of the experimental slice on the Allen Mouse CCF post linear transformation. (C) The registered CCF brain is overlaid on an experimental coronal slice. White frame indicates the chosen area from medial prefrontal cortex region which was extracted and visualized in (E,F). Green fluorescence signal was captured with 520nm excitation (E) and red fluorescence signal was captured with 640nm excitation wavelength (F).

### 2.4. Immunofluorescence staining

The BV2 cell sample grown on the glass-bottom dish were fixed with 4% paraformaldehyde and permeabilized with 0.1% Triton X-100 in PBS. Next, the sample was incubated with specific primary antibodies (Supplementary Table 1) and sencondary antibodies. Subsequently, the cell sample was preserved with antifade reagent and keep at 4°C in the dark till imaging. Fluorescent images were captured using Carl Zeiss LSM700 Confocal Microscope. Fluorescence signal intensity was analyzed using ImageJ Fuji software.

### 2.5. Digital PCR assay

EcoHIV proviral DNA was quantified by digital PCR using PSI, R5, and Pol Rainbow assay probes. Genomic DNA was extracted using the Zymo Quick-DNA Miniprep Kit (Zymo, Tustin, CA). Digital PCR was performed on the QIAcuity system (96-well format, 8.5K partitions per well; Qiagen, Germantown, MD). Quantification and data analysis were conducted using QIAcuity Software Suite (v1.2).

### 2.6. Statistical analysis

The independent samples *t*-test (SPSS Statistics 27, IBM Corp., Somer, NY), analysis of variance (ANOVA; SPSS Statistics 27), or regression statistical techniques (GraphPad Prism 5.02, GraphPad Software, Inc., La Jolla, CA) were selected for data analysis. The GraphPad Prism 5 software was chosen to create figures. An alpha level of *p*≤0.05 was considered as statistical significance.

## 3. Results

### 3.1. Increased GSDMD expression in EcoHIV-induced microglial pyroptosis in mice

Our previous study demonstrated that EcoHIV-infected animals exhibited neurocognitive impairments and synaptic dysfunction, accompanied by activation of the NogoA–NgR3/PirB–RhoA neuroinflammatory signaling pathway. Notably, microglia play a critical role as HIV-1 viral reservoirs. To better understand functional brain circuits, behavior, and neuropathology following HIV-1 infection, it is essential to characterize cell–cell connectivity, cellular density, and viral distribution across the entire brain. To address this challenge, we introduced a high-resolution volumetric whole-brain imaging approach using light-sheet fluorescence microscopy. Combined with NeuroInfo software, brain-wide cellular and molecular signals were automatically detected and quantitatively analyzed to assess HIV-1 viral distribution and GSDMD expression following EcoHIV infection. C57BL/6J wild-type mice received retro-orbital EcoHIV infusion and were analyzed at 0, 14-, 30-, 60-, and 90-days (d) post-infection. Whole brains were fixed, cleared using BINAREE solution, and incubated with specific primary antibodies. As shown in Figure 2A, whole-brain imaging using the BrightSLICE light-sheet microscope enabled visualization of EcoHIV distribution, with green fluorescence indicating viral infection. The medial prefrontal cortex (mPFC) was extracted to assess HIV-1 expression within a specific functional brain region (Figure 2B). Integrated fluorescence intensity was quantified using NeuroInfo software. As shown in Figure 2C, substantial HIV-1 signals were detected throughout the brain, even at 90 days post-infection. Surprisingly, viral expressions in the mPFC peaked at 14 days and remained elevated throughout the longitudinal study (Figure 2D). Our statistical analysis also presented EcoHIV viral distribution pattern in CA1 (Figure 2E), Dentate gyrus (Figure 2D) and whole hippocampus (Figure 2G) areas. GSDMD activation is a key marker of pyroptosis and neuroinflammation during HIV-1 infection. To evaluate pyroptotic activity, cleared whole-brain tissues were incubated with an anti-GSDMD primary antibody to assess temporal changes in expression following EcoHIV infection. Strikingly, GSDMD expression was markedly increased throughout the brain (Figure 2H-K). Compared with controls, mice at 14 days post-infection exhibited significantly elevated GSDMD levels, which remained high at 30 days post-infection. These findings highlight the pathological significance of pyroptosis in HIV-1–associated neuroinflammation.

**Figure 2.**
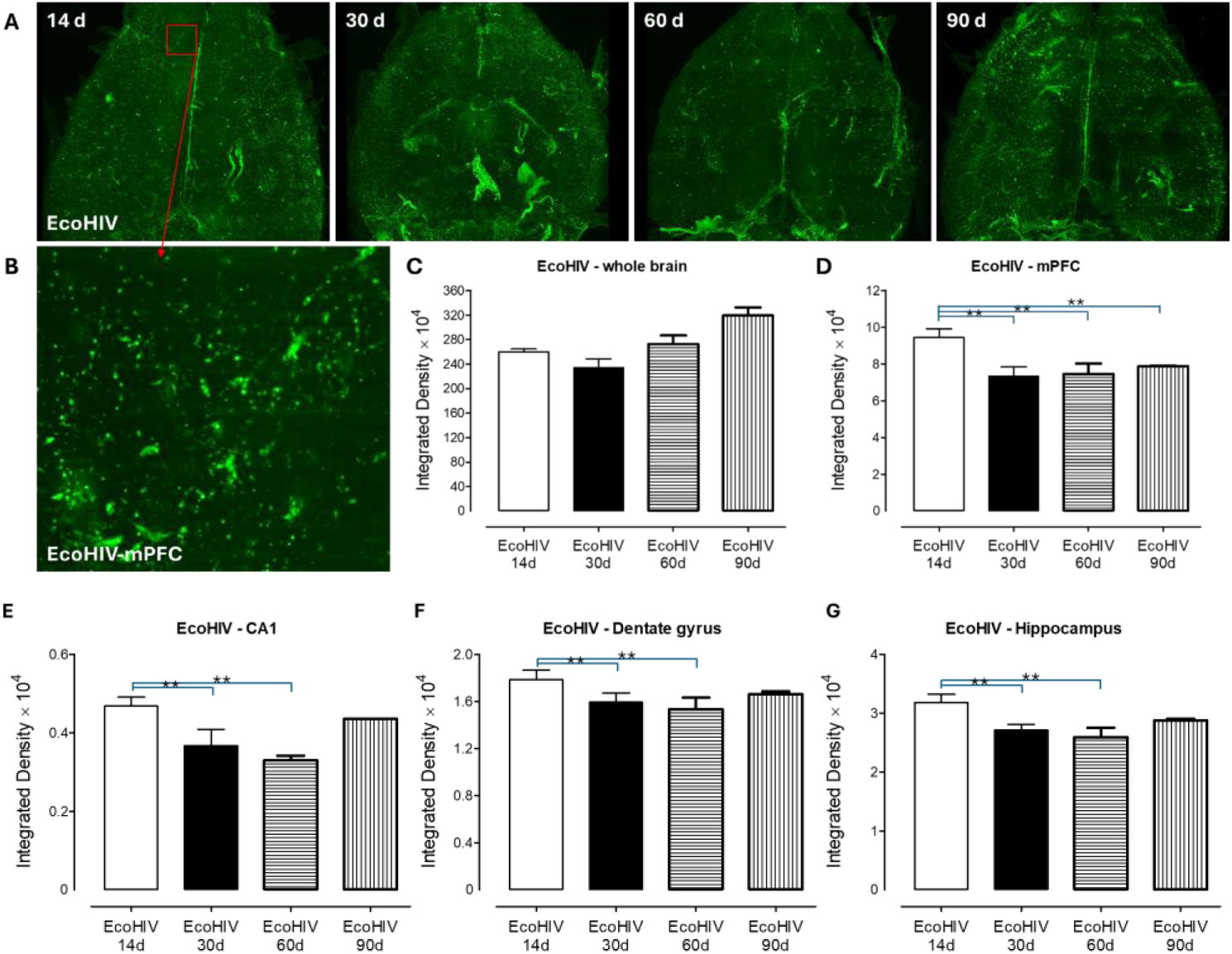
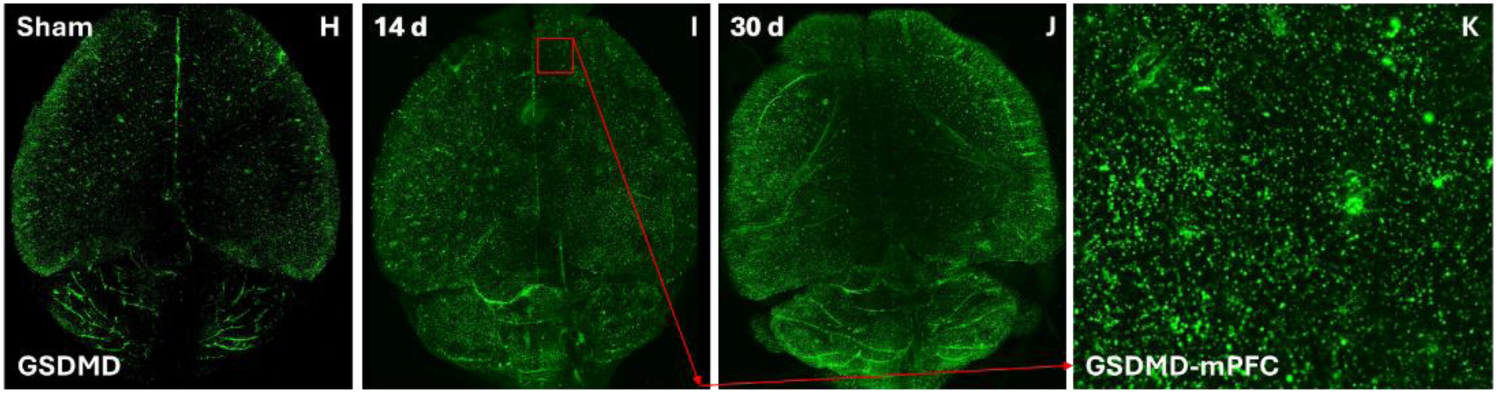
EcoHIV-eGFP lentivirus successfully induced HIV-1 infection and microglial pyroptosis in C57BL6/J mice. C57BL6/J wild type mice were randomly assigned into sham or EcoHIV infusion (at 14-, 30-, 60-, 90 days). The mouse brain tissue was fixed with 4% PFA and cleared by BINAREE kit. The anti-eGFP and anti-GSDMD primary antibody was utilized to assess the distribution of EcoHIV and microglia pyroptosis in mouse brain. The whole brain tissue imaging was taken by BrightSLICE light sheet microscope Imaging system using a 10× objective. (A) Representative images of eGFP signals at multiple times post EcoHIV infusion in mice captured with 520nm excitation. Positive expression indicates EcoHIV distribution. (B) Medial prefrontal cortex (mPFC) region (red frame in 2A) was extracted from the entire cleared brain. (C) Statistical assessment of integrated fluorescence density of EcoHIV-eGFP expression was performed in whole brain tissue at certain time points using NeuroInfo analysis software. (D) Statistical graph showed the monitored EcoHIV intensity in mPFC area based on NeuroInfo analysis. Bars indicate mean ± SEM. (E-G) Statistical analysis of EcoHIV distribution in CA1 (E), dentate gyrus (F), and whole hippocampus regions (G). (H-K) Representative images of GSDMD (a cellular marker for pyroptosis) at multiple times (0d, 14d, 30d) post the event and captured with 520nm excitation. Whole brain (H-J) and extracted mPFC (K) region (red frame in 2I) showing increased expression of GSDMD with a peak at 14 days after EcoHIV infection compared to control group.

### 3.2. The longitudinal association of HIV-1 and NLRP3 inflammasome during EcoHIV-induced microglial pyroptosis in mice

HIV-1 infection has been previously reported to activate myeloid cells leading to the secretion of proinflammatory factors, such as: IL-1β, TNFα, IL-6 [19, 20]. According to an *in vitr*o study, researchers exposed BV2 cells (a microglia cell line) under multi-doses of Tat (one of the HIV-1 toxic protein) and found an upregulated expression of microglial NLRP3 inflammasome in a dose-and time-dependent manner [21]. It was well-documented that GSDMD-mediated pyroptosis function through inflammasome activation, and NLRP3 inflammasome is the most prominent one during neuroinflammation [22]. To characterize pathological mechanism of microglial pyroptosis during HIV-1 infection, we subsequently analyzed NLRP3 inflammasome expression and distribution on cleared whole brain tissue which had been infused with EcoHIV for 0-, 14-, 30-, 60-, and 90 days. However, the detailed mechanism of microglia HIV-1 remained poorly understood. In the current study, we investigated whether exposure to EcoHIV increases NLRP3 inflammasome expression and mapped the distribution of its expression in entire brain tissue. As anticipated, EcoHIV-eGFP infusion for 14-, 30-, 60-, and 90 days significantly increased NLRP3 inflammasome expression in whole brain tissue, with a peak of 14 days and remained at an elevated level of expression post HIV-1 infection (Figure 3A), related to control group (0 day). Representative images of NLRP3 inflammasome expression were visualized through BrightSLICE light sheet microscope system. Statistical analysis by NeuroInfo software, as shown in Figure 3C, revealed NLRP3 activation after HIV-1 infection at 14 days (292.190 ± 26.508), 30 days (259.415 ± 23.485), 60 days (252.551 ± 8.985) and 90 days (225.894 ± 20.403). Statistical assessment of the extracted mPFC region (Figure 3B&D) suggested a similar trend of NLRP3 inflammasome expression in Figure 3C. We also evaluated NLRP3 expressions in CA1 (3E), dentate gyrus (3F), and hippocampus regions (3G). The next step was to examine the potential association between microglial HIV-1 infection and expression of NLRP3 inflammasome. As shown in Figure 3H-L, we observed a much higher proportion of colocalization of NLRP3 inflammasome expression in EcoHIV^+^ cells in whole brain, suggesting a critical pathology characterized by the NLRP3 inflammasome in microglia during HIV-1 associated neuroinflammation. Collectively, these data imply that HIV-1 prime the NLRP3 inflammasome via enhancing NLRP3 expression in mouse brain, the latter further results in microglial pyroptosis.

**Figure 3.**
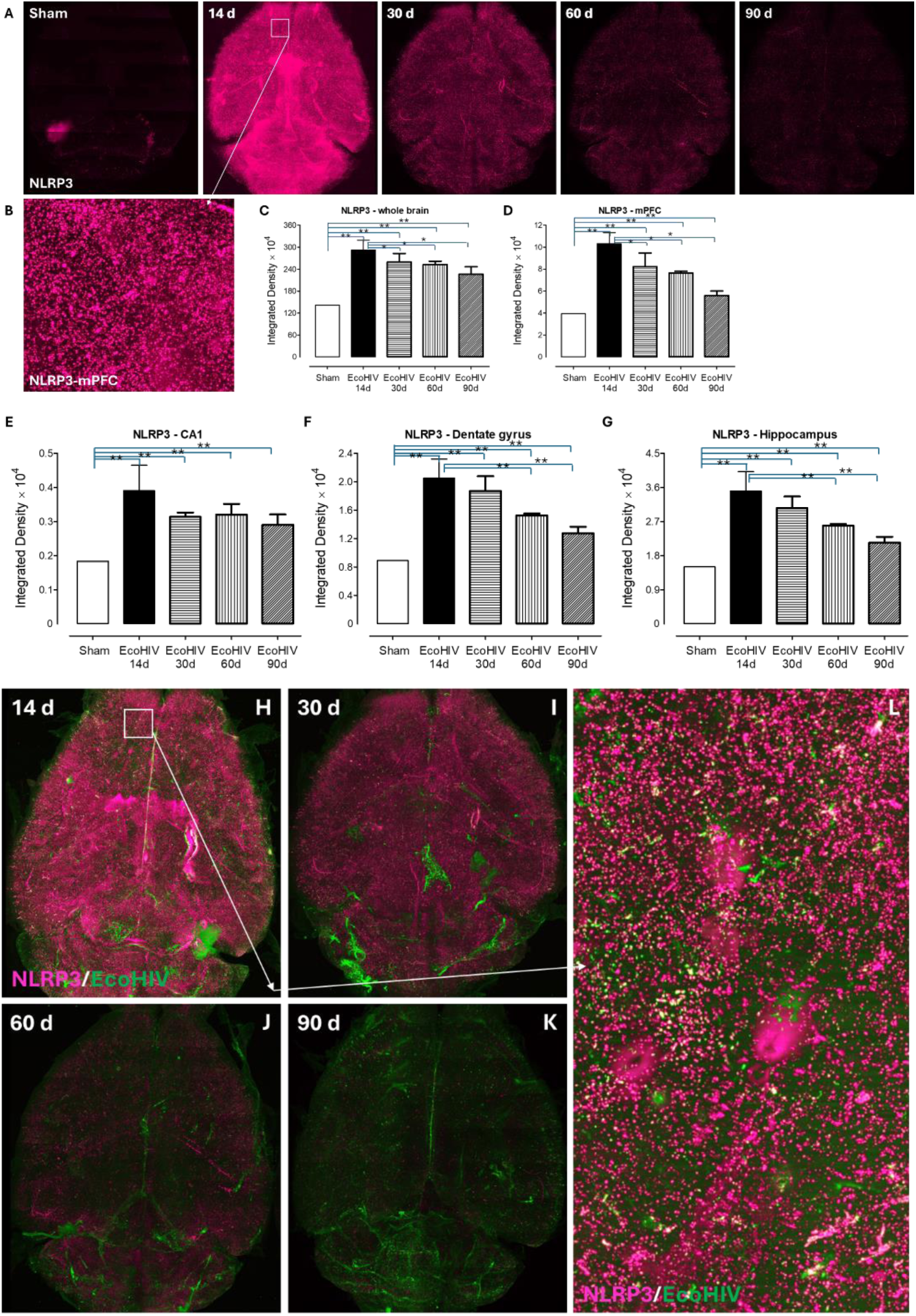
EcoHIV-induced microglial pyroptosis was associated with NLRP3 inflammasome activation in mice. C57BL6/J wild type mice were randomly assigned into sham or EcoHIV infusion (at 14d, 30d, 60d, 90 days). The mouse brain tissue was fixed with 4% PFA and cleared by BINAREE kit. Anti-NLRP3 primary antibody was used to monitor the alterations of neuroinflammation after EcoHIV infection. The whole brain tissue imaging was taken by BrightSLICE light sheet microscope Imaging system using a 10× objective. (A,B) Representative images of NLRP3 inflammasome at multiple times (0d, 14d, 30d, 60d, 90 days) after EcoHIV infection and was captured with 640nm excitation. Whole brain (A) and extracted mPFC (B) region (white frame in 3A) displaying increased expression of NLRP3 at different times post EcoHIV infusion with a peak at 14 days. (C, D) Statistical analysis described NLRP3 inflammasome expression in whole brain (C) and mPFC (D) area at different times post EcoHIV. Bars indicate mean ± SEM. Statistical significance was assessed using student t test and p-values were presented. * Stand for P<0.05 and ** indicated P<0.01. (E-G) Statistical evaluation of NLRP3 distribution in CA1 (E), dentate gyrus (F), and whole hippocampus (G) regions. (H-K) Representative images showing the colocalization of NLRP3^+^ signals in EcoHIV-infected eGFP^+^ cells in whole brain at 14d (H), 30d (I), 60d (J) and 90d (K) after EcoHIV infusion. White frame in 3H indicates representative HIV/NLRP3 colocalization in the chosen area from mPFC area which was extracted and visualized in (L).

### 3.3. Pharmacological inhibition of NLRP3 alleviates EcoHIV-induced microglial pyroptosis and immune exhaustion in vitro

Having demonstrated EcoHIV infection upregulated the microglial NLRP3 inflammasome, we then investigated whether the NLRP3 inhibitor could alleviate EcoHIV-induced microglial neuroinflammation. We hypothesized that inhibition of NLRP3 inflammasome would decrease GSDMD expression which further attenuated microglial pyroptosis and its immune response to the HIV-1 infection. To address this, BV2 cells (a mouse microglia cell line) were exposed to EcoHIV lentivirus for 3 days following the treatment of MCC950 (NLRP3 inhibitor, 10μM), and analyzed for GSDMD, NLRP3 and Caspase1 expression. The viral expression in EcoHIV-infected BV2 microglia was detected by using digital PCR and measured as (7.4 ± 0.62) × 10^4^ cp/µL and (3.7 ± 0.311) × 10^6^ cp/µg DNA. Meanwhile, MCC950 treatment did not alternate EcoHIV expression in BV2 cells (data not shown). Our results showed that EcoHIV infection induced an increase in total amount of GSDMD and NLRP3 expression in BV2 cells (including both EGFP^+^ and EGFP^-^ cells, see attached Supplementary Figure 1). Remarkably, as presented in Figure 4A, we observed a statistically significant decrease in GSDMD expression (10.23 ± 4.97 vs. 23.37 ± 3.33) in EcoHIV^+^ microglia (EGFP^+^) after MCC950 treatment compared to EcoHIV only (Figure 4J). NLRP3 inflammasome activation involves a two-step process: enhancing levels of NLRP3 and triggering the processing of Caspase1 which subsequently leads to the release of IL-1β [23, 24]. We next quantified NLRP3 and Caspase1 in EcoHIV^+^ BV2 cells; the data in Figure 4D&G suggested that a significant change was found in EcoHIV^+^ microglia between the groups of EcoHIV infected cells with or without NLRP3 inhibitor. Both NLRP3 (13.95 ± 2.20 vs. 24.58 ± 3.63) and Caspase1 (46.60 ± 7.14 vs. 59.15 ± 12.21) expressions in EcoHIV^+^ microglia were significantly reduced after MCC950 treatment related to EcoHIV infection group w/o MCC950 (Figure 4K&L). However, MCC950 itself did not alter GSDMD, NLRP3 and Caspase1 expression in BV2 cells. These data highlight the importance of NLRP3 inflammasome activation on microglial pyroptosis during HIV-1 associated neuroinflammation.

**Figure 4.**
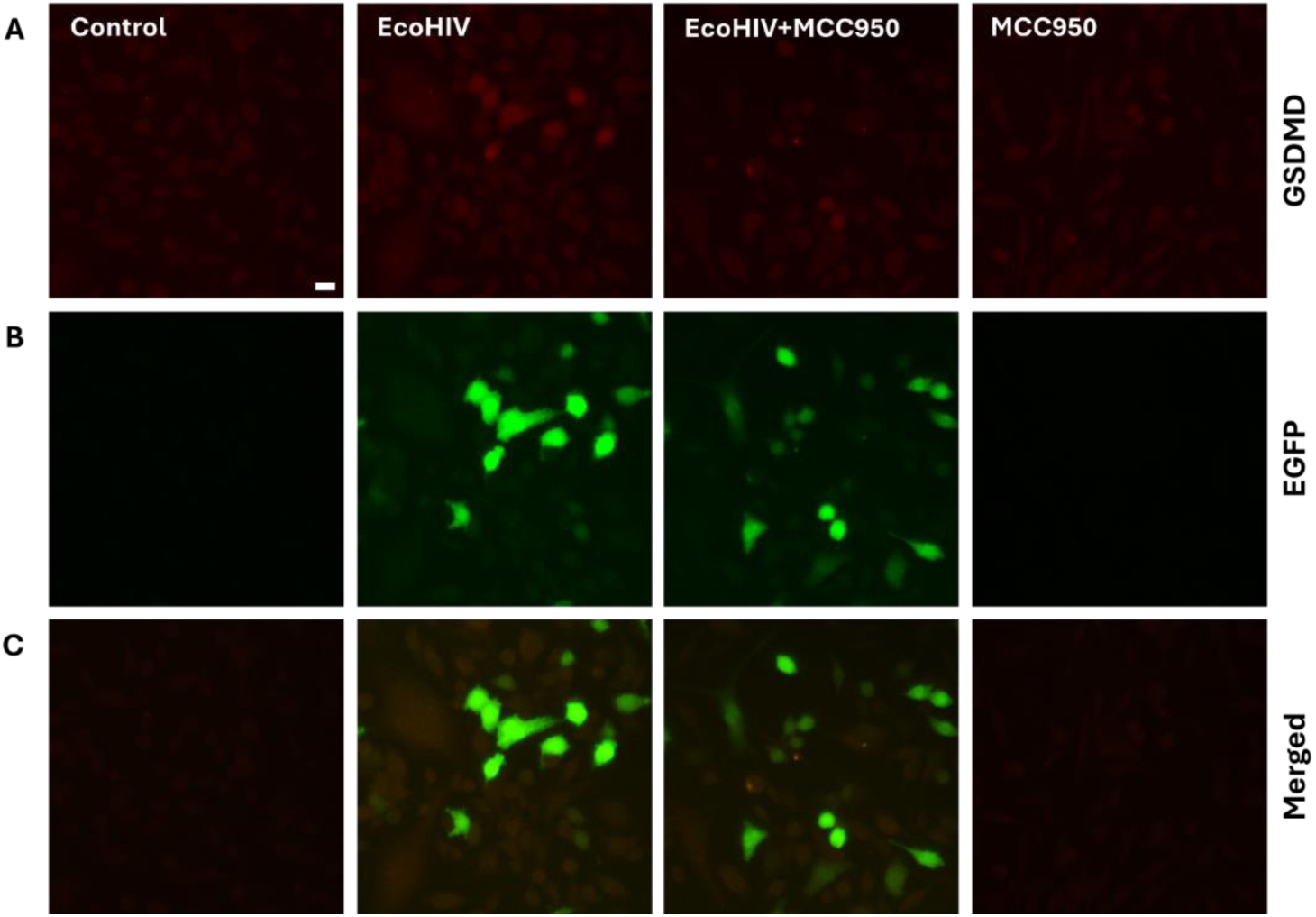
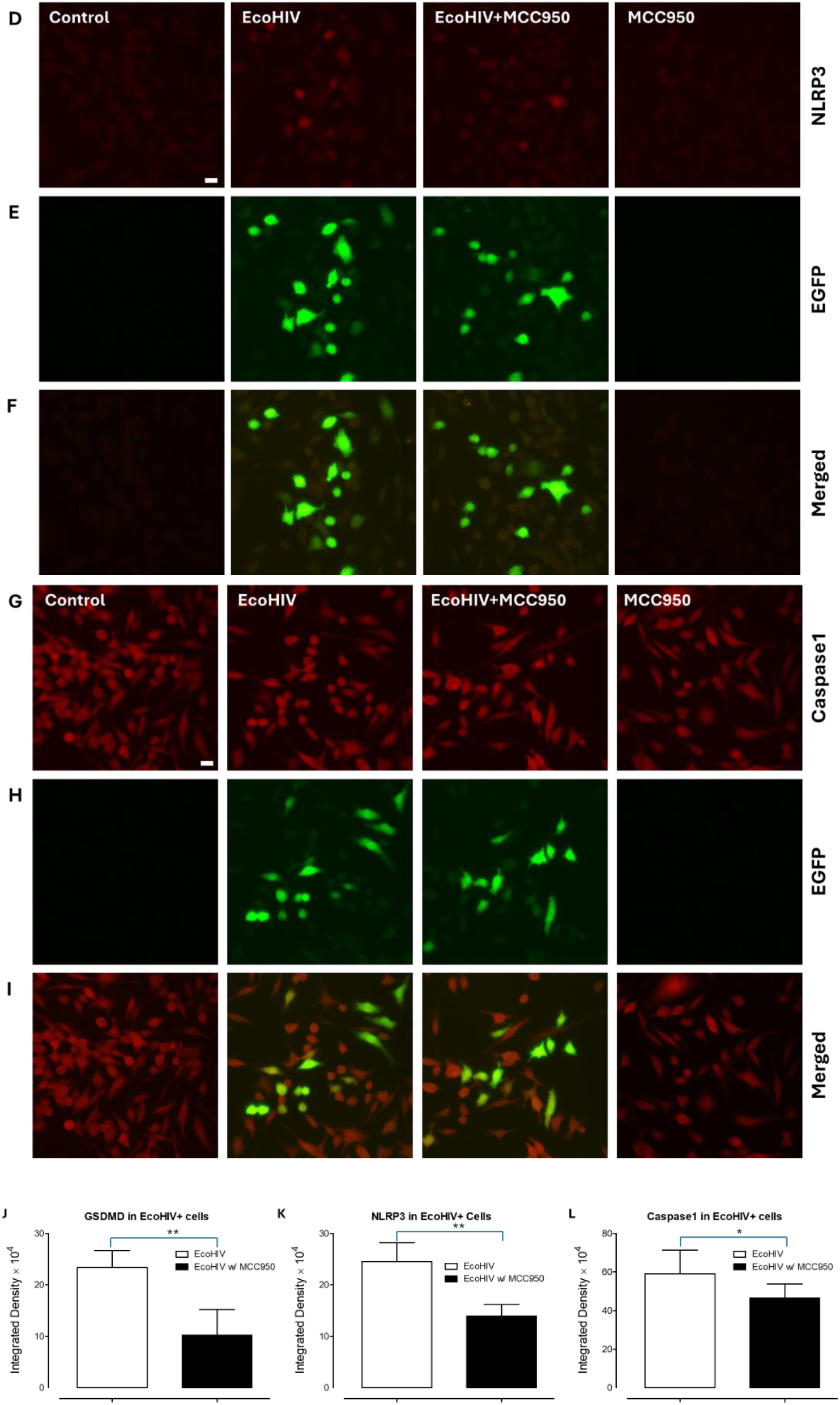
NLRP3 inflammasome inhibitors attenuates microglial pyroptosis in EcoHIV-infected BV2 cells. (A-C) Representative confocal images showing fluorescence signals of GSDMD^+^ (red, A-C), NLRP3^+^ (red, D-F) and Caspase1^+^ (red, G-I) in control and EcoHIV infected BV2 microglia cells w or w/o MCC950 treatment. The green fluorescence signals of eGFP signify the EcoHIV-infected microglia in BV2 cell line *in vitro*. The dual-labeling of EcoHIV^+^/GSDMD^+^ (C), EcoHIV^+^/NLRP3^+^ (F) and EcoHIV^+^/Caspase1^+^ (I) show the colocalization of targeted proteins in EcoHIV-infected cells. (J-L) Quantification of integrative density of HLA-DR (J), PD-1 (K) and Ki67 (L) expression in immunofluorescence-stained EcoHIV^+^ cells, classified by HIV and MCC950 status. Data are presented as mean ± SEM; p-values determined by *t*-test are indicated. Scale bar, 50 µm.

To determine whether NLRP3 inflammasome activation is functionally involved in microglial immune exhaustion, we assessed several markers of cellular immunity, including HLA-DR, PD-1, and Ki67. HLA-DR class II molecules of the human major histocompatibility complex (MHC) present processed extracellular antigens to CD4+ helper T lymphocytes. It is widely accepted that HLA-DR is characterized as a state of immune cellular activation [25, 26]. Microglia upregulate HLA-DR in response to interferon (IFN)-γ stimulation and in different pathological conditions, including MS [27, 28, 29]. Programmed cell death protein 1 (PD-1) is an immune checkpoint receptor expressed on immune cells, including microglia, and functions to maintain immune homeostasis and prevent excessive immune activation and autoimmunity [30]. Ki67 is a nuclear protein strongly associated with cellular proliferation and has been linked to increased immune checkpoint expression. Although Ki67 is widely used as a proliferation marker, its role as a biomarker reflecting microglial immune responses during neuroinflammation remains to be fully elucidated [31]. As shown in Figure 5, treatment with an NLRP3 inflammasome inhibitor significantly attenuated the expression of HLA-DR, PD-1, and Ki67 in EcoHIV^+^ microglia; however, only moderate alteration was observed in the total BV2 cell population (including both eGFP⁺ and eGFP⁻ cells; Supplementary Figure 1). Then, we quantified HLA-DR, PD-1 and Ki67 expression specifically in EcoHIV⁺ microglia. As shown in Figure 4J-K, significant differences were observed between EcoHIV^+^ cells treated with or without MCC950. Both HLA-DR (25.68 ± 2.54 vs. 59.29 ± 5.07) and PD-1 (12.09 ± 3.53 vs. 23.55 ± 5.69) and Ki67 (18.39 ± 4.93 vs. 63.08 ± 4.69) expression levels were significantly reduced in EcoHIV⁺ microglia following MCC950 treatment compared with the EcoHIV infection group without MCC950 (Figure 4K and L). In contrast, MCC950 alone did not alter the expression of GSDMD, NLRP3, or caspase-1 in BV2 cells. Collectively, these data suggest that activation of the microglial NLRP3 inflammasome contributes to microglial immune exhaustion during HIV-1 infection, and that pharmacological inhibition of NLRP3 may improve microglial immune dysfunction and alleviate microglial pyroptosis during HIV-1–associated neuroinflammation.

**Figure 5.**
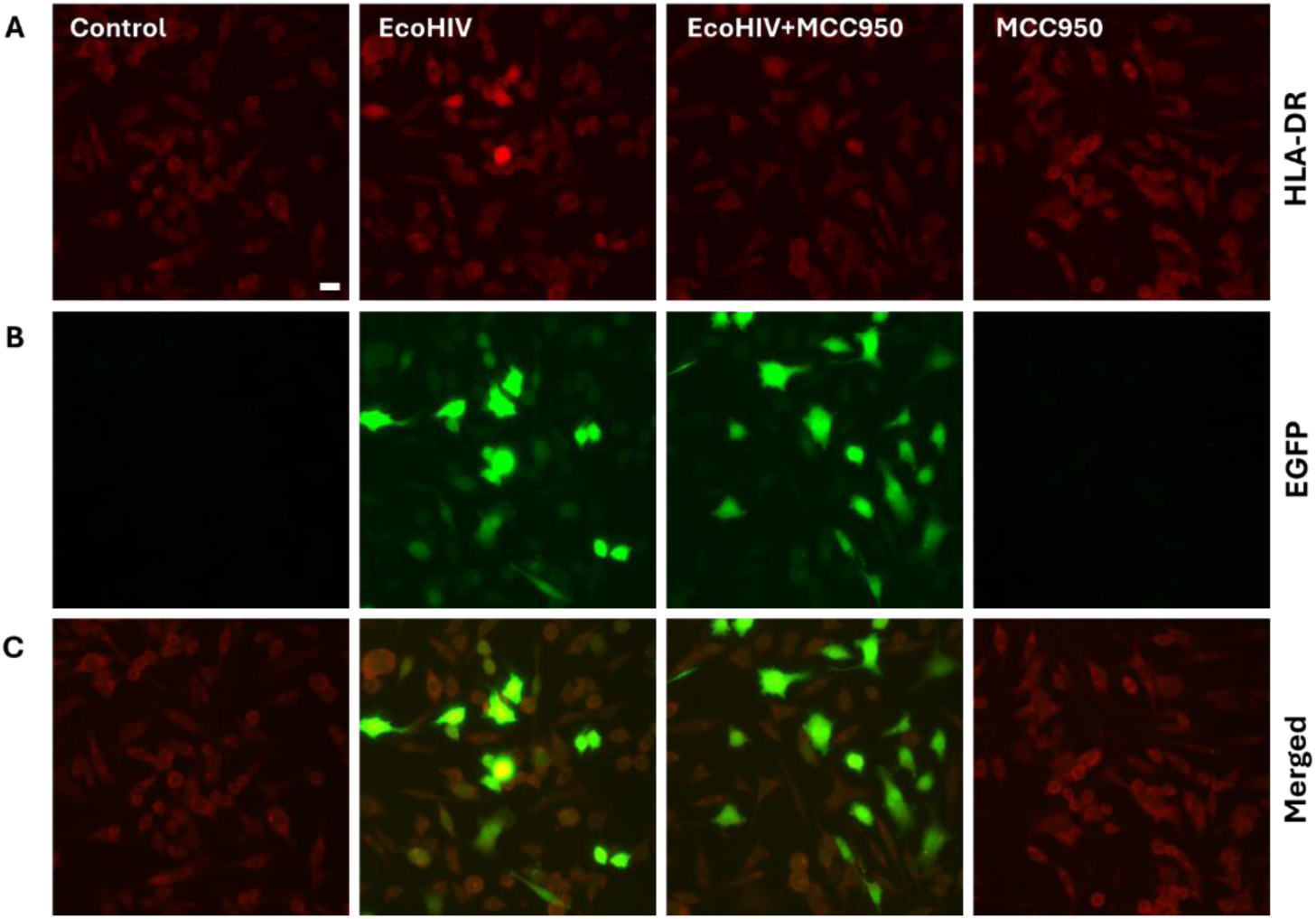
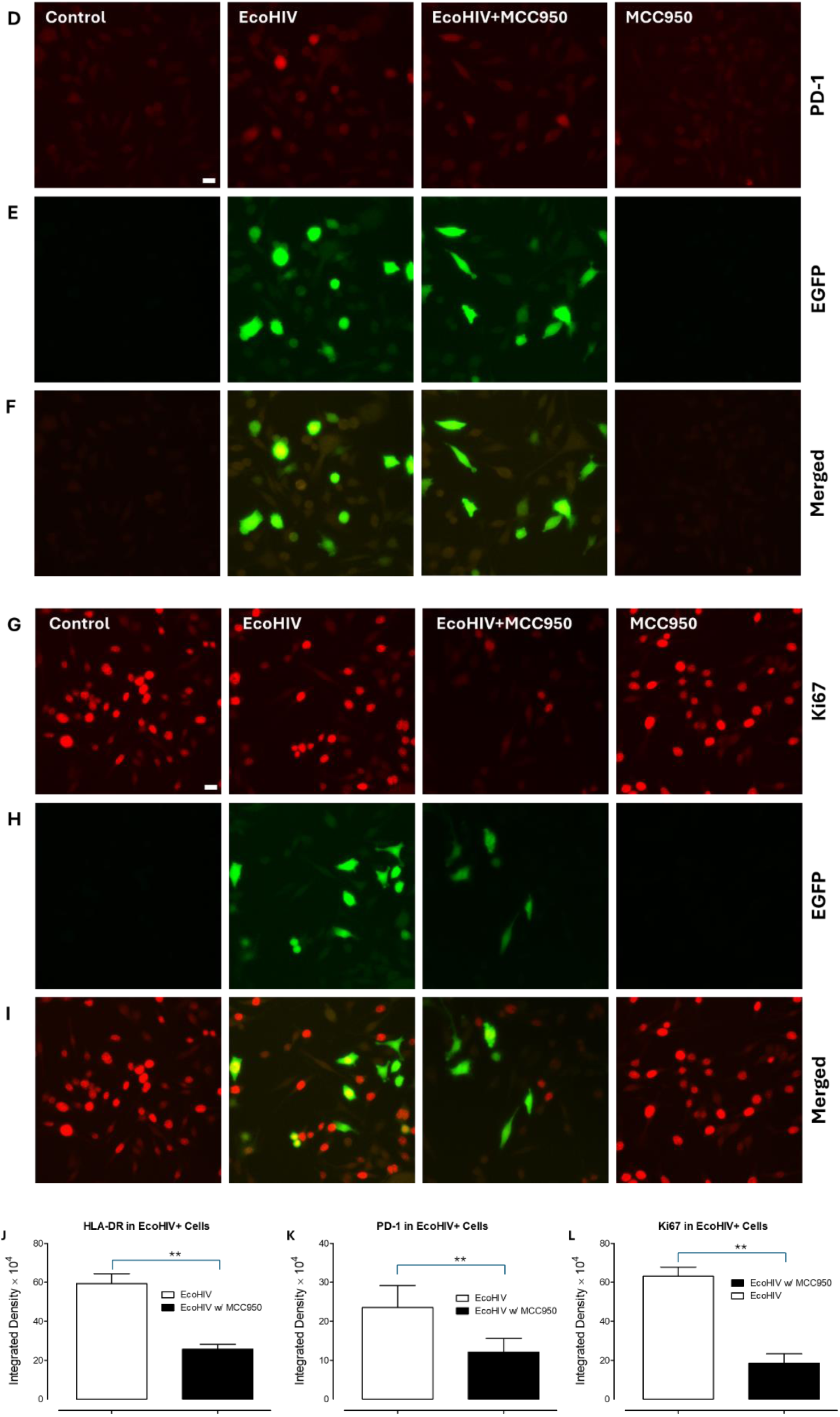
NLRP3 inflammasome inhibitor attenuates microglial immune exhaustion in HIV-infected BV2 cells. (A-C) Representative confocal images showing fluorescence signals of HLA-DR^+^ (red, A-C), PD-1^+^ (red, D-F) and Ki67^+^ (red, G-I) in control and EcoHIV-infected BV2 microglia cells with or without MCC950 treatment. The green fluorescence signals of eGFP stand for the EcoHIV-infected microglia in BV2 cell line *in vitro*. The dual-labeling of EcoHIV^+^/HLA-DR^+^ (C), EcoHIV^+^/PD-1^+^ (F) and EcoHIV^+^/KI67^+^ (I) show the colocalization of targeted proteins in EcoHIV-infected cells. (J-L) Quantification of integrative density of HLA-DR (J), PD-1 (K) and Ki67 (L) expression in immunofluorescence-stained EcoHIV^+^ cells, classified by HIV and MCC950 status. Data are presented as mean ± SEM; p-values determined by *t*-test are indicated. Scale bar, 50 µm.

We further analyzed the proportional changes of GSDMD, NLRP3, Caspase1, HLA-DR, PD-1 and Ki67 in EcoHIV^+^ microglia w/ or w/o MCC950 interference. Remarkably, we observed that NLRP3 inflammasome inhibition utilizing MCC950 significantly decreased the percentage of double positive cell numbers of GSDMD^+^/EcoHIV^+^ (20.42% ± 7.37% vs. 73.96% ± 8.91%, P < 0.01), NLRP3^+^/EcoHIV^+^, HLA-DR^+^/EcoHIV^+^ (37.75% ± 7.16% vs. 76.40% ± 5.29%, P < 0.01), PD-1^+^/EcoHIV^+^ (48.27% ± 8.01% vs. 76.03% ± 8.45%, P < 0.01) and Ki67^+^/EcoHIV^+^ (48.61% ± 3.98% vs. 79.79% ± 3.73%, P < 0.01) related to EcoHIV^+^ microglia without MCC950 treatment, suggesting the NLRP3 inhibitor improved microglial immune exhaustion and decreased NLRP3-associated microglial pyroptosis.

Overall, these findings indicated that HIV-1-induced NLRP3 inflammasome activation in microglia, the latter sequentially leading to microglial pyroptosis status during HIV-1 infection.

## 4. Discussion

Despite the use of cART, however, there is still an increase in prevalence of asymptomatic neurocognitive impairment (ANI) or mild neurocognitive disorder (MND) in cART-treated patients which account for up to 70% of HAND [32, 33]. Substantial evidence suggests that HIV-1 protein-induced neuroinflammation (such as via Tat, gp120 etc.) contribute a critical role in the progression of HAND. One of the studies has reported that HIV-1 Tat protein was detected in the brains of infected individuals and also implicated pyroptosis in the progressions of HIV-1 pathogenesis [34]. In the present study, we investigated the occurrence and association of NLRP3 inflammasome activation with microglia pyroptosis both in EcoHIV-infected C57BL6/J mice and the BV2 microglial cell line *in vitro*. Remarkably, we were able to perform whole-brain detection and analysis of HIV-1 distribution and NLRP3 inflammasome expression through the “SLICE” light-sheet microscope/NeuroInfo system. Our data showed a significant EcoHIV infection in the rodent brain as early as 14 days after viral infusion, predominantly in medial prefrontal cortex area and mainly localized in microglia. The persistent HIV-1 viral expression was assessed at 30, 60, and 90 days after infection. Moreover, we identified the NLRP3 inflammasome as a sensor that drives cell pyroptosis in EcoHIV-infected microglia. From our *in vitro* study, MCC-950 (NLRP3 inflammasome inhibitor) blocked microglia pyroptosis characterized by decreasing GSDMD, NLRP3, and Caspase1 in EcoHIV+ cells and improved microglia immune dysfunction in which several hall markers of immune exhaustion, such as: HLA-DR, PD-1 and Ki76 were altered by MCC-950 treatment. Based on these findings, a proposed mechanism of HIV-1 induced NLRP3-associated microglial pyroptosis is portrayed in the schematic in Fig. 6. These findings not only further our understanding of the HIV-1 pathogenesis but also elucidate potential innovative strategies for host immune homeostasis in HIV-1 infection.

**Figure 6.**
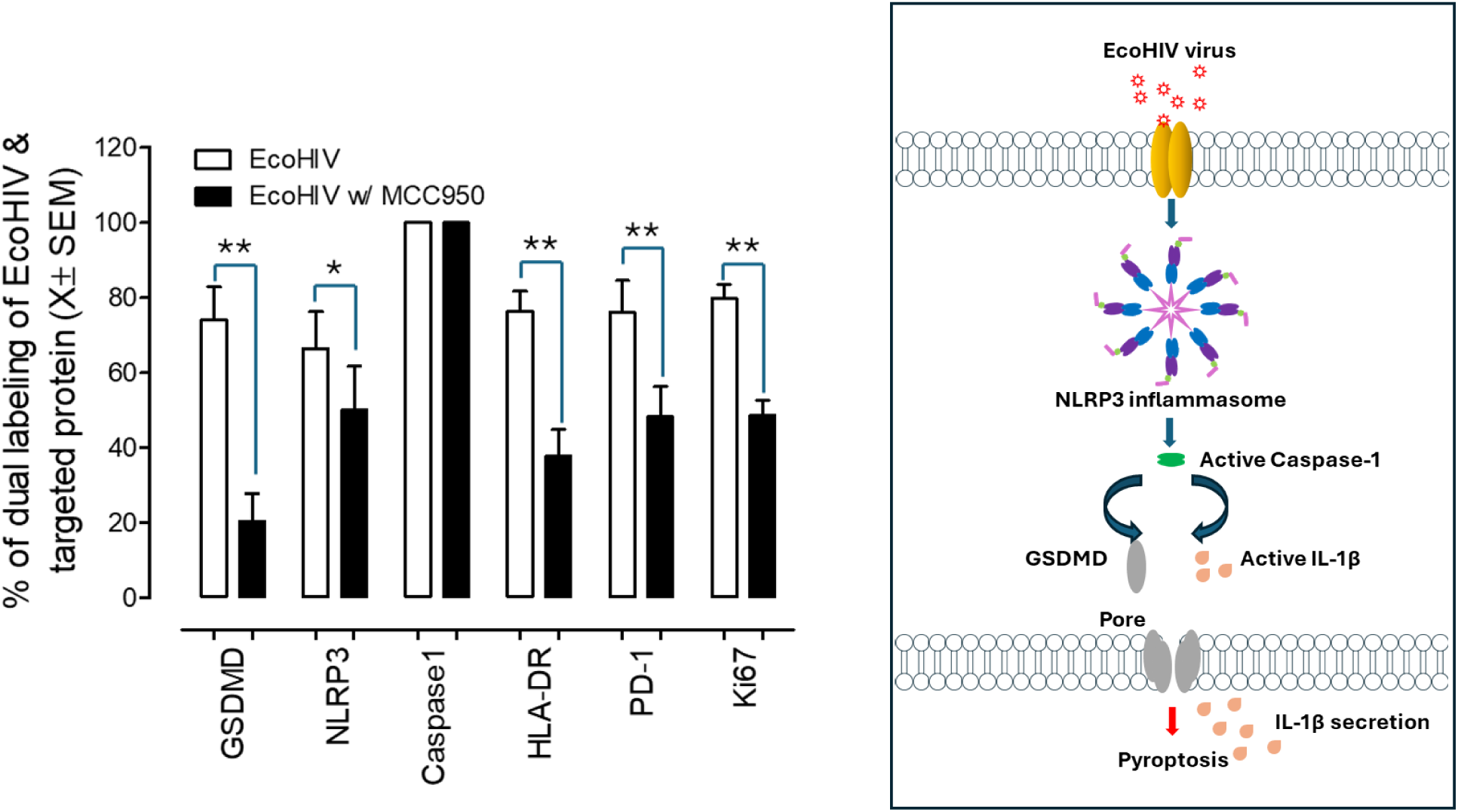
NLRP3 inflammasome inhibitor (MCC950) alleviated EcoHIV-related microglial pyroptosis and prevented microglial immune exhaustion in EcoHIV-infected BV2 cells *in vitro*. Quantification of integrative density of GSDMD, NLRP3, Caspase1, HLA-DR, PD-1 and Ki67 expression in immunofluorescence-stained cell samples, classified by HIV and MCC950 status. Data are presented as mean ± SEM; p-values determined by *t*-test are indicated. * Stands for P<0.05 and ** indicates P<0.01. Schematic maps show the signal pathway of NLRP3 inflammasome associated microglia pyroptosis after EcoHIV infection.

Characterizing NLRP3 inflammasome-associated microglial pyroptosis and its spatial distribution across the entire rodent brain is essential for understanding functional brain circuits and their relationships to complex behavior and pathology. However, whole-brain analysis poses substantial technical challenges. In addition, whole-brain imaging generates massive datasets that complicate data stitching, visualization, and analysis. Even after successful data acquisition and assembly, a further challenge remains: the lack of simple, accurate, and standardized methods for quantifying and comparing cell populations across the entire brain. Here, we introduce an end-to-end platform – BrightSLICE microscope/NeuroInfo system for imaging and analyzing intact, optically cleared rodent brains that substantially lowers the barrier to whole-brain analysis and enables robust, brain-wide cellular quantification. Our findings show that substantial HIV-1 expressions from entire brain area are apparent at 14, 30, 60, 90 days after EcoHIV-eGFP infusion. Meanwhile, one of the hallmarks of cellular pyroptosis – GSDMD was markedly upregulated throughout the brain, remained consistently high at 30 days post infection. Utilizing the BrightSLICE light sheet microscope in combination with the NeuroInfo system, we successfully assessed the quantification of GSDMD expression and HIV-1 distribution in the current study. It has been well documented that EcoHIV predominantly infects microglia in brain of rodent models [35]. Now we have presented evidence of microglial pyroptosis in the progression of HIV-1.

Pyroptosis is characterized as an inflammatory programmed cell death process and has a pivotal role in various neurological diseases [36]. Pyroptosis is also distinguished with its unique morphological and pathophysiological features, including chromatin condensation, DNA fragmentation, inflammatory caspases activation and secretion of proinflammatory cytokines [37, 38]. The induction of pyroptosis is finely regulated to precisely defend against pathogenic invaders and maintain homeostasis of the host. Additionally, pyroptosis is distinct from apoptosis characterized as simultaneously induced cell death and triggering a cascade of inflammatory responses. Among the GSDM family, GSDMD is the most extensively studied protein and functions as an important executor of inflammasome-associated pyroptosis, leading to disease-related inflammation [39]. In this progression, NLRP3 inflammasome activation predominantly induced procaspase1 assembly and resulting Caspase1-dependent pyroptosis and pore formation. The NLRP3 inflammasome is a multiprotein complex composed of Nod-like receptors, apoptosis-associated speck-like proteins containing a caspase recruitment domain and caspase1 precursor. Its activation involves two steps: First, a range of stimulates bind with Nod-like receptors and forming a ASC complex to activate caspase1; Then, activated caspase1 cleaves the pore forming protein gasdermin D (GSDMD) and secretion of IL-1β and IL-18, resulting in pyroptosis [40].

Recent research has implicated cellular pyroptosis participation in the progression of HAND, evidenced as gp120-induced microglial NLRP3 inflammasome activation resulting in pyroptosis and IL-1β release [41] Additionally, inflammasome activation has been found in microglia in Alzheimer’s disease, in which the AD-associated neuropathogenic proteins stimulate the NLRP3 inflammasome signaling pathway and induced caspase1 activation, resulting in microglial polarization to a proinflammatory phenotype [42, 43]. However, Miao et al. [44] found that HIV-infected individuals who abuse methamphetamine exhibit microglial pyroptosis via activating AIM2 inflammasome.

In many neurodegenerative disorders, as a resident immune macrophage in the CNS, microglia initially perform as an anti-inflammatory phenotype, followed by a fast switch to a predominate proinflammatory phenotype, which modulates neuroinflammation and homeostasis in the CNS. In addition, we observed elevated colocalization of NLRP3 inflammasome and HIV eGFP in microglia in the entire brain region, especially in media prefrontal cortex. Therefore, BV2 cells were employed as a study subject for *in vitro* experiments to elucidate the mechanisms of NLRP3 inflammasome-mediated microglial pyroptosis during HIV. Similarly, in our study, using MCC950 to alleviate NLRP3 inflammasome activation in BV2 cells significantly reduced the level of pyroptosis and increased cell viability. Collectively, these findings suggest that METH and HIV-1 Tat proteins can synergistically induce microglial pyroptosis through dsDNA damage-mediated activation of the AIM2 inflammasome.

In a long-term outpatient cohort study, researchers explored the relationship between the HIV reservoir and CD8⁺ T-cell activation. Notably, persistently elevated HLA-DR expression on CD8⁺ T cells was observed and was positively correlated with total HIV DNA levels [45, 46]. These findings suggest that the latent HIV reservoir may contribute to a sustained inflammatory state in people living with HIV despite long-term viral suppression. Consistent with this, previous longitudinal studies have demonstrated that the expression of HLA-DR and PD-1 on CD8⁺ T cells closely correlate with HIV DNA levels in both peripheral blood and lymph nodes [47, 48].

HLA-DR, a member of the MHC class II molecule family, is widely recognized as a hallmark of chronic immune activation, reflecting continuous antigen production and presentation of HIV DNA reservoir [49]. PD-1 and ki67 are also characterized as important receptors to achieve a more comprehensive understanding of HIV-1 associated microglia immune homeostasis. PD-1 is a central regulator of T cell exhaustion and contributes to maintenance of the latent HIV infection. Previous studies reported an upregulation of PD-1 on HIV-specific CD4 T cells, and its expression was correlated with HIV viremia [50, 51, 52]. In addition, blockade of the PD-1 activation with an anti-PD-L1 antibody showed restoration of HIV CD4 T cell proliferation [53]. Our previous work demonstrated enhanced microglial proliferation in the medial prefrontal cortex of EcoHIV rats, accounting for 55.9% of the variance in synaptophysin intensity. Specifically, an increased number of co-localized Iba1^+^ and Ki67^+^ cells was significantly associated with lower synaptophysin intensity, its role in the immune dysregulation associated with HIV infection [54]. In the current study, attenuation of NLRP inflammasome activation using MCC950 significantly reduced the expression levels of HLA-DR, PD-1 and Ki67 (over two folds compared to EcoHIV group without MCC950) in microglia *in vitro*, with reductions exceeding two-fold compared to the EcoHIV group without MCC950 treatment. We therefore hypothesized that modulation of NLRP3 inflammasome activation in microglia may alleviate HIV induced microglia pyroptosis, potentially through NLRP3/caspase1/GSDMD signaling axis. Our hypothesis is supported by recent findings demonstrating upregulated transcription of NLRP3 and caspase-1 in HIV-1 infected patients exhibiting poor immune recovery [55].

Nevertheless, several limitations should be acknowledged. First, the predominant infected cell type in the EcoHIV mouse model is microglia, which does not fully recapitulate the cellular landscape of HIV-1 infection in humans. Despite this limitation, the model remains valuable for investigating microglial function in HIV-1-associated neurodegeneration. Future studies incorporating postmortem brain tissue from HIV-infected individuals receiving combination antiretroviral therapy (cART) will be essential to achieve a more comprehensive understanding. Second, the potential neuroprotective effects of NLRP3 inhibition by MCC950 should be further validated in EcoHIV mice to substantiate targeting microglial pyroptosis as a therapeutic strategy for HIV-1-associated neurocognitive disorders (HAND).

In summary, this study provides evidence that NLRP3-dependent microglial pyroptosis may represent a dominant pathway contributing to HIV-1 disease progression in the central nervous system. Targeting the NLRP3/caspase-1/GSDMD/IL-1β axis may therefore offer novel therapeutic opportunities to improve HIV-1-associated neurocognitive and affective outcomes.

## Supplementary Materials

Yes.

## Ethics approval and consent to participate

All procedures were approved by the Institutional Animal Care and Use Committee (IACUC) of the University of South Carolina (Federal Assurance #D16-00028).

## Authors’ Contributions

HL designed and conducted all the experiments and was a major contributor in writing the manuscript. CM and RB provided full support on experimental design and edited and revised the manuscript. MS provided advice on manuscript revision. All authors read and approved of the final manuscript.

## Informed Consent Statement

None.

## Availability of data and materials

All data are available in the main text or supplementary file.

## Acknowledgments

None.

