## Supplemental data table 1 & fIgure 1 for "NLRP3 inflammasome-related microglial pyroptosis in EcoHIV infected mice"

**Table 1. List of primary antibodies**

| Antigen | Host Species | Type | Company | Catalogue Number | Antibody ID |
|---|---|---|---|---|---|
| **Anti-GSDMD antibody** | Rabbit | Monoclonal | Abcam | **AB219800** | **AB_219800** |
| **Anti-NLRP3 antibody** | Rabbit | Monoclonal | Abcam | **AB270449** | **AB_270449** |
| **Anti-pro Caspase-1 + p10 + p12 antibody** | Rabbit | Monoclonal | Abcam | **AB179515** | **AB_179515** |
| **Goat Anti-Rabbit IgG H&L (Alexa Fluor® 647)** | Goat | Monoclonal | Abcam | **AB150083** | **AB_150083** |
| **Goat Anti-Mouse IgG H&L (Alexa Fluor® 647)** | Rabbit | Monoclonal | Abcam | **AB150115** | **AB_150115** |
| **Goat Anti-Rabbit IgG H&L (Alexa Fluor® 555)** | Rabbit | Monoclonal | Abcam | **AB150078** | **AB_150078** |
| **Goat Anti-Mouse IgG H&L (Alexa Fluor® 555)** | Rabbit | Monoclonal | Abcam | **AB150114** | **AB_150114** |
| **Ki67** | Mouse | Monoclonal | Abcam | **AB279653** | **AB_279653** |
| **Anti-GFP antibody [9F9.F9]** | Mouse | Monoclonal | Abcam | **AB1218-1003** | **AB_1218** |

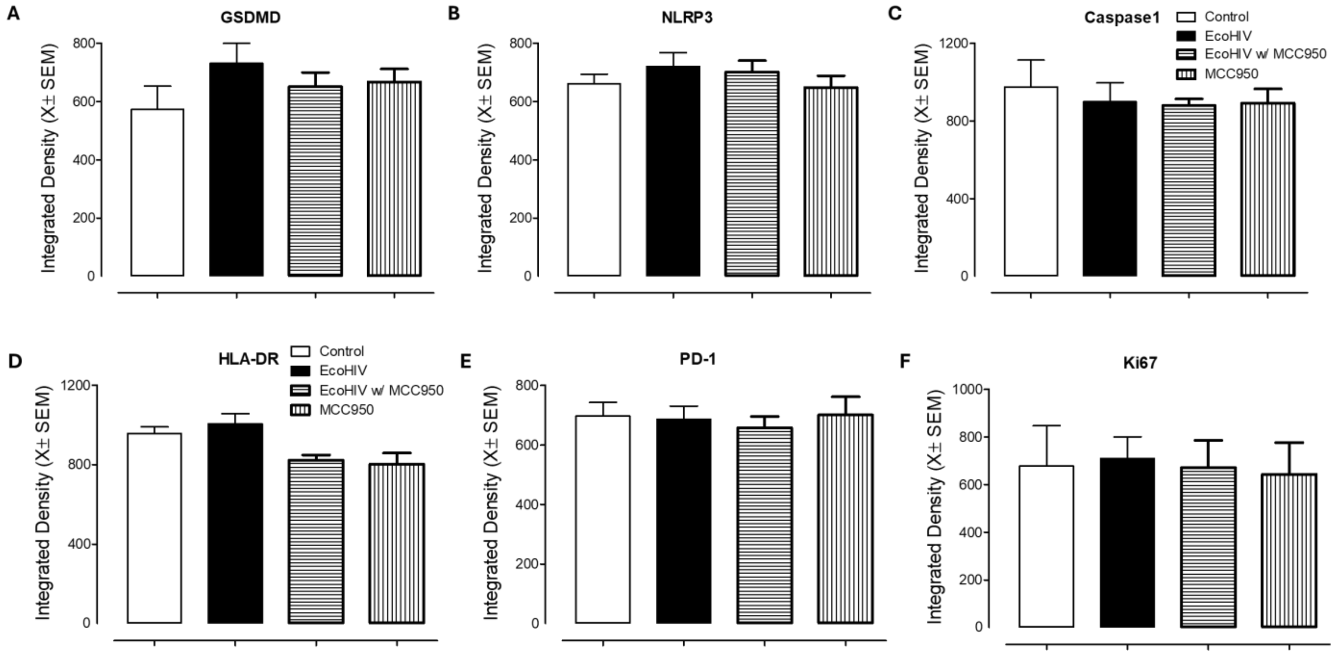


**Supplementary Figure 1. NLRP3 inflammasome inhibitors attenuates microglial pyroptosis in EcoHIV-infected BV2 cells.** Quantification of integrative density of GSDMD (A), NLRP3 (B), Caspase1 (C), HLA-DR (D), PD-1 (E) and Ki67 (F) expression in the overall experimental BV2 cells, classified by HIV and MCC950 status.
